# Identification of oncolytic vaccinia restriction factors in canine high-grade mammary tumor cells using single-cell transcriptomics

**DOI:** 10.1101/2020.05.29.123133

**Authors:** Béatrice Cambien, Kevin Lebrigand, Alberto Baeri, Nicolas Nottet, Catherine Compin, Audrey Lamit, Olivier Ferraris, Christophe N Peyrefitte, Virginie Magnone, Jérôme Henriques, Laure-Emmanuelle Zaragosi, Sophie Giorgetti-Peraldi, Frédéric Bost, Marine Gautier-Isola, Roger Rezzonico, Pascal Barbry, Robert Barthel, Bernard Mari, Georges Vassaux

## Abstract

Mammary carcinoma, including triple-negative breast carcinomas (TNBC) are tumor-types for which human and canine pathologies are closely related at the molecular level. Low-passage, primary carcinoma cells from TNBC versus non-TNBC were used to compare the efficacy of an oncolytic vaccinia virus (VV). We show that non-TNBC cells are 28 times more sensitive to VV than TNBC cells in which VV replication is impaired. Single-cell RNA-seq performed on two different TNBC cell samples infected or not with VV highlighted three distinct populations: naïve cells, bystander cells, defined as cells exposed to the virus but not infected and infected cells. The transcriptome of these three populations showed striking variations in the modulation of pathways regulated by cytokines and growth factors. We hypothesized that the pool of genes expressed in the bystander populations was enriched in antiviral genes. Bio-informatic analysis suggested that the reduced activity of the virus was associated with a higher mesenchymal status of the cells. In addition, we demonstrate experimentally that high expression of one gene, DDIT4, is detrimental to VV production. Considering that DDIT4 is associated with a poor prognosis in various cancers including TNBC, out data highlight DDIT4 as a candidate resistance marker for oncolytic poxvirus therapy. This information could be used to design new generations of oncolytic poxviruses. Beyond the field of gene therapy, this study demonstrate that single-cell transcriptomics can be used to identify cellular factors influencing viral replication.

**Author summary:** The identification of cellular genes influencing viral replication/propagation have been studied using hypothesis-driven approaches and/or high-throughput RNA interference screens. In the present report, we propose a methodology based on single-cell transcriptomic. We have studied, in the context of oncolytic virothepary, the susceptibility of primary, low-passage mammary carcinoma cells of canine origin from different grades to an oncolytic vaccinia virus (VV). We highlight a fault in replication of VV in cells originated from high-grade triple-negative breast carcinomas (TNBC). Single-cell RNA-seq performed on TNBC cell samples infected with VV suggested that the reduced activity of the virus was associated with a higher mesenchymal status of the cells. We also demonstrate that high expression of one gene, DDIT4, is detrimental to VV production. Considering that DDIT4 is associated with a poor prognosis in various cancers including TNBC, out data highlight DDIT4 as a candidate resistance marker for oncolytic poxvirus therapy. Beyond the field of cancer gene therapy, we demonstrate here that single-cell transcriptomics increases the arsenal of tools available to identify cellular factors influencing viral replication.

## Introduction

Oncolytic vaccinia virus (VACV) represent a new class of anticancer agents with multiple mechanisms of action. VACV has been shown to act at three distinct levels [1]. VACV infects and selectively replicates in cancer cells, leading to primary oncolysis and resulting in cancer cell destruction [2]. It also disrupts the tumor vasculature [3], and reduces tumor perfusion. Finally, the release of tumor antigens from dead tumor cells participates to the initiation of an immune response that may be effective against tumor cells [1, 4–7]. In humans, VACV, administered intratumourally or systemically has been well-tolerated in various clinical trials [4].

Poxviruses are large viruses with cytoplasmic sites of replication and are considered less dependent on host cell functions than other DNA viruses. Nevertheless, the existence of cellular proteins capable of inhibiting or enhancing poxvirus replication and spread has been demonstrated. Cellular proteins such as dual specific phosphatase 1 DUSP1 [8] or barrier to autointegration factor (BAF) [9] have been shown to be detrimental to the virus. In contrast, the ubiquitin ligase cullin-3 has been shown to be required for the initiation of viral DNA replication [10]. Furthermore, high-throughput RNA interference screens have suggested the potential role of hundreds of proteins acting as either restricting or promoting factors for poxviruses [10–13]. These studies highlight the importance of cellular factors in VACV replication and spread. Theoretically, over-expression or down-regulation of these putative restricting or promoting factors in carcinoma cells could result in reduced sensitivity and even resistance to VACV when primary oncolysis is considered. The concept of resistance to primary oncolysis by VACV has, so far, not been formally demonstrated. For example, in the field of breast cancer research, *in vitro* testing in established human cell lines and *in vivo* xenografts in mice, have shown clearly and convincingly that VACV has anti-tumor activity against breast cancer [14, 15]. The efficacy of VACV was evident in triple-negative high-grade breast carcinoma mouse models [15], a pathology associated with poor prognosis and for which new therapeutic options are urgently needed. Nevertheless, these studies were performed in established cancer cell lines that may differ from the actual carcinoma cells present in the tumors.

Spontaneously occurring mammary cancers in dogs are of potential interest in the development of new anticancer agents [16–18] as the classification of canine breast carcinoma is relevant to that of human’s [19–23]. If differences have been highlighted in complex carcinomas [24], simple canine carcinomas faithfully represent human breast carcinomas, both at the histological and molecular level [20, 21]. This is particularly the case for the so called “triple negative carcinomas” (lack of estrogen and progesterone receptors and of epidermal growth factor receptor type 2) [22, 25, 26], for which therapeutic options are limited and unsatisfactory.

The aim of the present study was to determine whether differences in VACV-induced ability to kill freshly-isolated primary cells from low-grade versus high-grade canine breast carcinomas could be demonstrated. Bulk and single cell RNA-seq were used to analyze events associated with VACV infection and to characterize genes potentially interfering with the VACV cycle.

## Results

### TNBC canine cells show reduced sensitivity to a vaccinia virus-Lister strain deleted in the thymidine kinase gene (VV) compared to TNBC carcinoma cells

Cells from TNBC or non-TNBC were infected with vaccinia virus-Lister strain deleted in the thymidine kinase gene (VV) at different multiplicities of infection (MOI) and the numbers of cells remaining in the culture wells were monitored after four days. Figure 1A presents an example of dose-response curves showing that non-TNBC cells are more sensitive to VV-mediated cell lysis than TNBC cells. This observation is in sharp contrast to the situation observed in human established cell lines in which MCF7 cells (as a representative of non-TNBC cells) and MDA-MB-231 cells (as a representative of TNBC cells) show an equivalent sensitivity to VV-induced cell lysis (Figure 1B). The combined LD50 of experiments performed on all available samples (n= 19 non-TNBC and n = 6 for TNBC) are presented in Figure 1C and show that non-TNBC cells are 28 times more sensitive to VV than TNBC cells. The viral production was compared in canine TNBC and non-TNBC cells. Quantitative PCR to determine the number of viral genomes produced upon infection (Fig. 1D) and titration to determine the number of viral particles showed a reduced number of infectious viral particles produced upon infection of TNBC cells (Fig. 1E).

**Figure 1:**
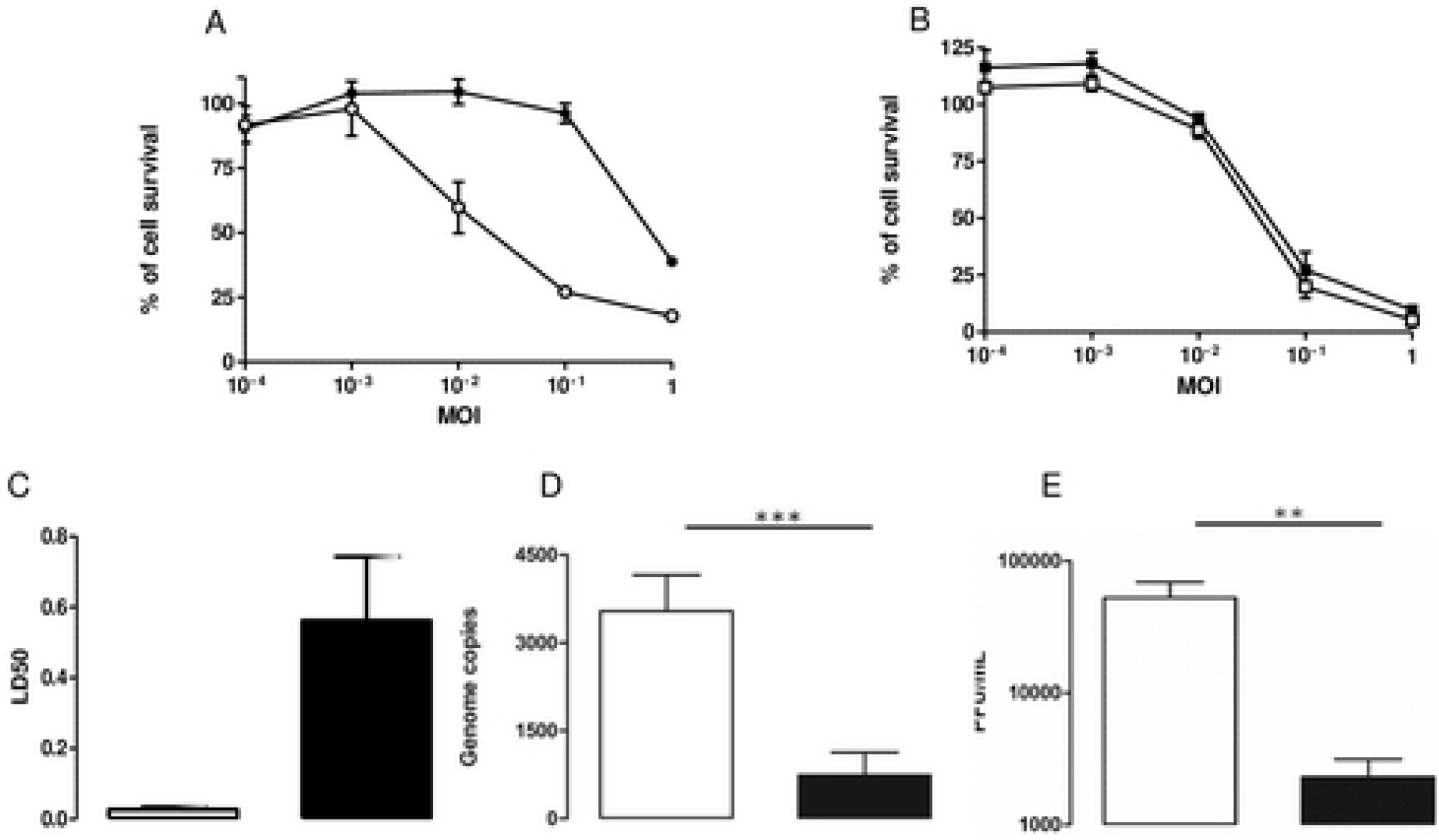
Comparison of the efficacy of VV on non-TNBC or TNBC of different origins. **A.** Primary canine cells (non TNBC: white circles; TNBC black circles) or **B.** human established cell lines (MCF7: white squares, as a non-TNBC cell line; MDA-MB-231: black squares, as a TNBC cell line) cells were infected at different MOIs with VV. Fours days later, the remaining cells were estimated using a MTT assay. The results are presented as a percentage of cell-survival in uninfected cells and are mean +/− SEM of six different experimental points. A similar experiment was performed on the whole cohort of cells and he LD50 (dose of virus capable of killing 50% of the cell population) was calculated. Data show the mean LD50 +/− SE obtained on 18 independent non-TNBC and 6 independent TNBC cultures (**C**). Twenty-four hours after VV infection, non-TNBC (white bars) or TNBC (black bars) cells were collected, DNA was isolated and subjected to quantitative PCR to titer the number of viral genome per well (**D**). Three days after VV infection, non-TNBC (white bars) or TNBC (black bars) cells were collected, homogenized and the number of infectious particles generated in the culture dishes were titrated (**E**).

### Replication as opposed to viral infection/early stage of viral transcription is affected mainly in canine TNBC cells

Infection and early-stages of viral transcription of a vaccinia virus-Copenhagen strain recombinant in which GFP expression is driven by an immediate-early vaccinia virus promoter was examined on TNBC or non-TNBC cells (Figure 2 A-D). Counting the number of propidium iodide-positive and GFP-positive cells revealed a statistically-significant, 5 % difference in infection/early-stage of viral transcription between the two types of cells (Fig. 1E). Propidium iodide staining showed a classical nuclear labelling as well as cytosolic dots (Figure 2 B and D). These structures are usually found in cells infected with VV and are often referred to as DNA factories or mininuclei [27, 28]. They are cytoplasmic sites of viral DNA replication [27]. The percentage of mininuclei-positive cells in GFP-positive cells 8 hours after infection was 53% and 26.6% in non-TNBC and TNBC cells, respectively (Fig. 1F). In mininuclei-positive cells, an average of 3 and 1 mininuclei were found in the cytosol of non-TNBC and TNBC cells, respectively (Fig.1G). Altogether, these data show that although a difference in the efficacy of infection/early stage of viral transcription can be detected in non-TNC and TNBC cells, the main quantitative difference lies in the number of DNA factories, as in an average population, 6 times more viral DNA factories can be detected in non-TNBC compared to TNBC cells.

**Figure 2:**
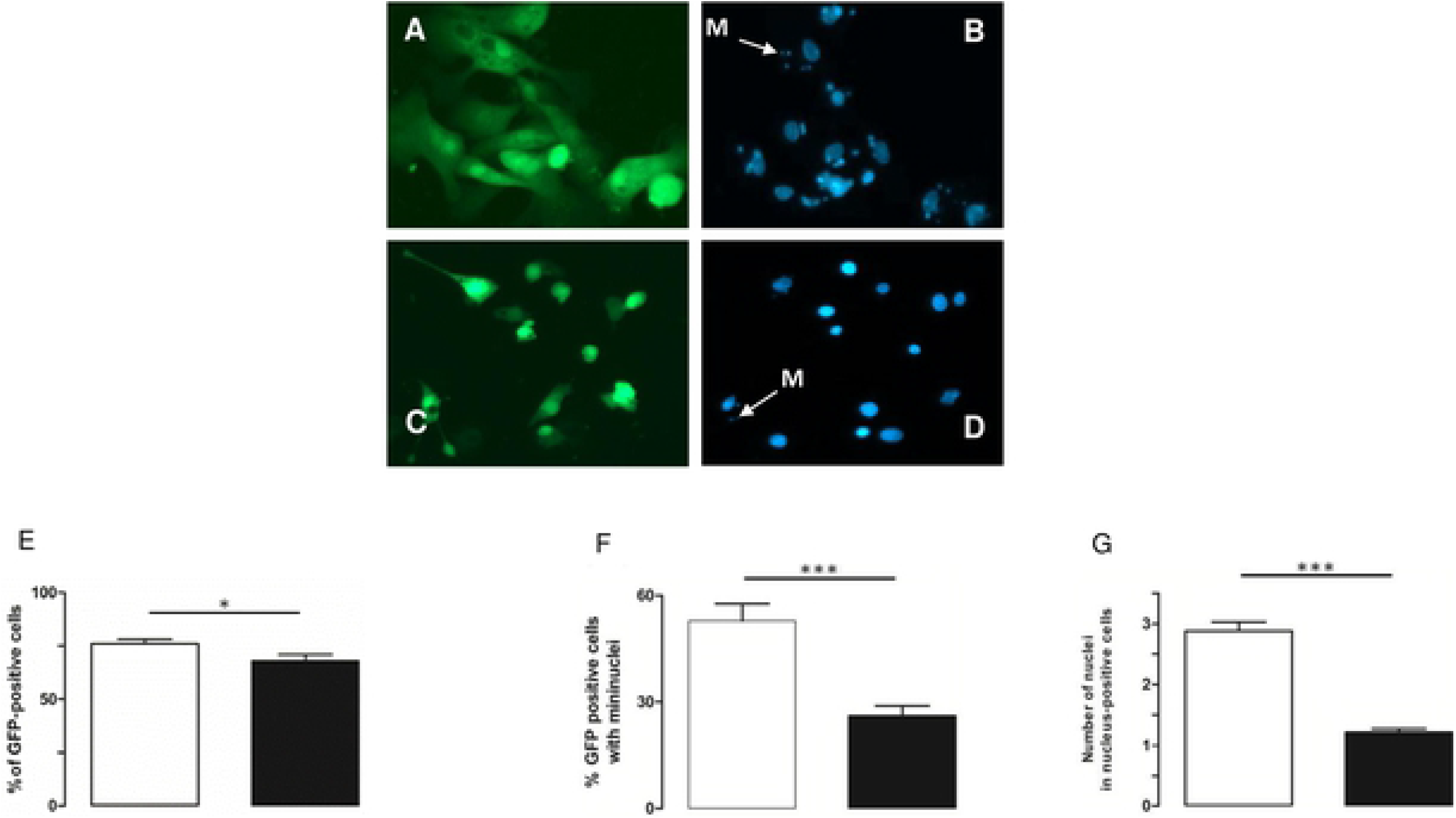
Primary canine TNBC cells infected with a VV virus exhibit a reduced numbers of mininuclei compared to primary canine non-TNBC cells. Non-TNBC (**A**, **B**) or TNBC (**C**, **D**) cells were infected with a VV in which the expression of the GFP is driven by an immediate-early VV promoter (MOI=5). Three hours after infection, the cells were fixed and stained with propidium iodide (PI). A and B: GFP staining; C and D: propidium iodide staining; M: mininuclei. The images presented are representative of 150 images obtained from 5 primary canine non-TNBC and TNBC. The number of PI and GFP positive cells was determined. The percentage of GFP+ cells (**E**), of mini-nuclei in GFP+ cells (**F**) and the number of mini-nuclei in nuclei-positive cells (**G**) are presented. (***: p < 0.001; ** p < 0.01; * p < 0.05).

Upon infection, early viral genes are rapidly transcribed by the viral RNA-polymerase packaged within the infectious particles [29]. By contrast, the expression of intermediate- and late-viral genes requires de novo protein synthesis and viral replication in DNA factories [29]. An implication of the results presented in Fig. 1 and 2 is that the expression of the early viral genes should be comparable in non-TNBC and TNBC, while the expression of intermediate and late genes should be largely impaired in TNBC. To assess this hypothesis, kinetics of expression of the early gene E9L and late gene A27L were performed on non-TNBC and TNBC. Figure S1 shows that, the expression of E9L is comparable in non-TNBC and TNBC cells 2 and 4 hours after infection and a difference in E9L expression is clearly visible 8 hours after infection. The late viral gene A27L was hardly detectable 2 and 4 hours after infection and its expression markedly increased 8h post-infection in non-TNBC, while A27L expression remained low at this time-point in TNBC. To extend these data, the kinetics of bulk viral RNA-expression in non-TNBC versus TNBC cells infected with VV was determined using another pair of donors. Figure 3 shows that the difference of expression of early viral genes in non-TNBC versus TNBC cells is detectable and is statistically significant. However, this difference is much greater when intermediate and late viral gene expression is concerned. Altogether, these data suggest that, although infection/very viral early gene expression is statistically-significantly lower in TNBC than in non-TNBC cells, the number of mininuclei, the replication of the virus and subsequent expression of the intermediate and late viral gene are quantitatively much more affected.

**Figure 3:**
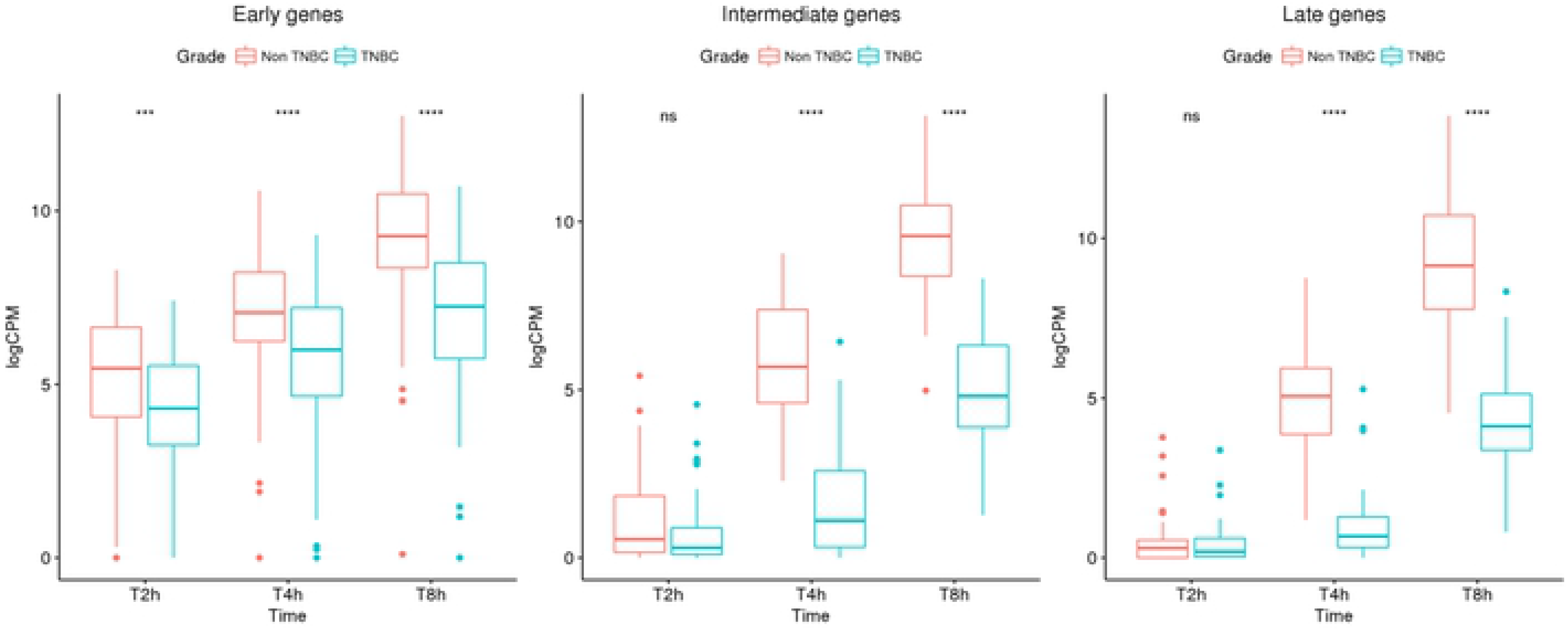
Comparison of the viral gene transcription in non-TNBC and TNBC carcinoma cells infected with VV. TNBC (blue) or non-TNBC (red) carcinoma cells were infected at a multiplicity of infection of 5. At different times post-infection (2, 4, 8 hours), the cells were collected and processed for RNA sequencing. The data represent the levels of early (left panel), intermediate (middle panel) or late (right panel) viral gene expression.

### Single-cell RNA sequencing to dissect VV infection of TNBC cells: impact of the infection on cellular genes

To characterize further the infection of TNBC cells by VV, we performed single-cell transcriptomic analysis. In these experiments, two independent TNBC primary cell cultures were either mock infected or infected with VV at a MOI of 5. Six hours later, the cells were trypsinized and subjected to the 10X Genomics single-cell protocol, followed by sequencing. Figure 4 shows that, in the two experiments performed, the number of cellular genes expressed decreases as the extent of viral gene expression increases. Furthermore, this decrease in the number of cellular genes expressed is more drastic in the subset of cells expressing higher levels of late viral genes (in red).

**Figure 4:**
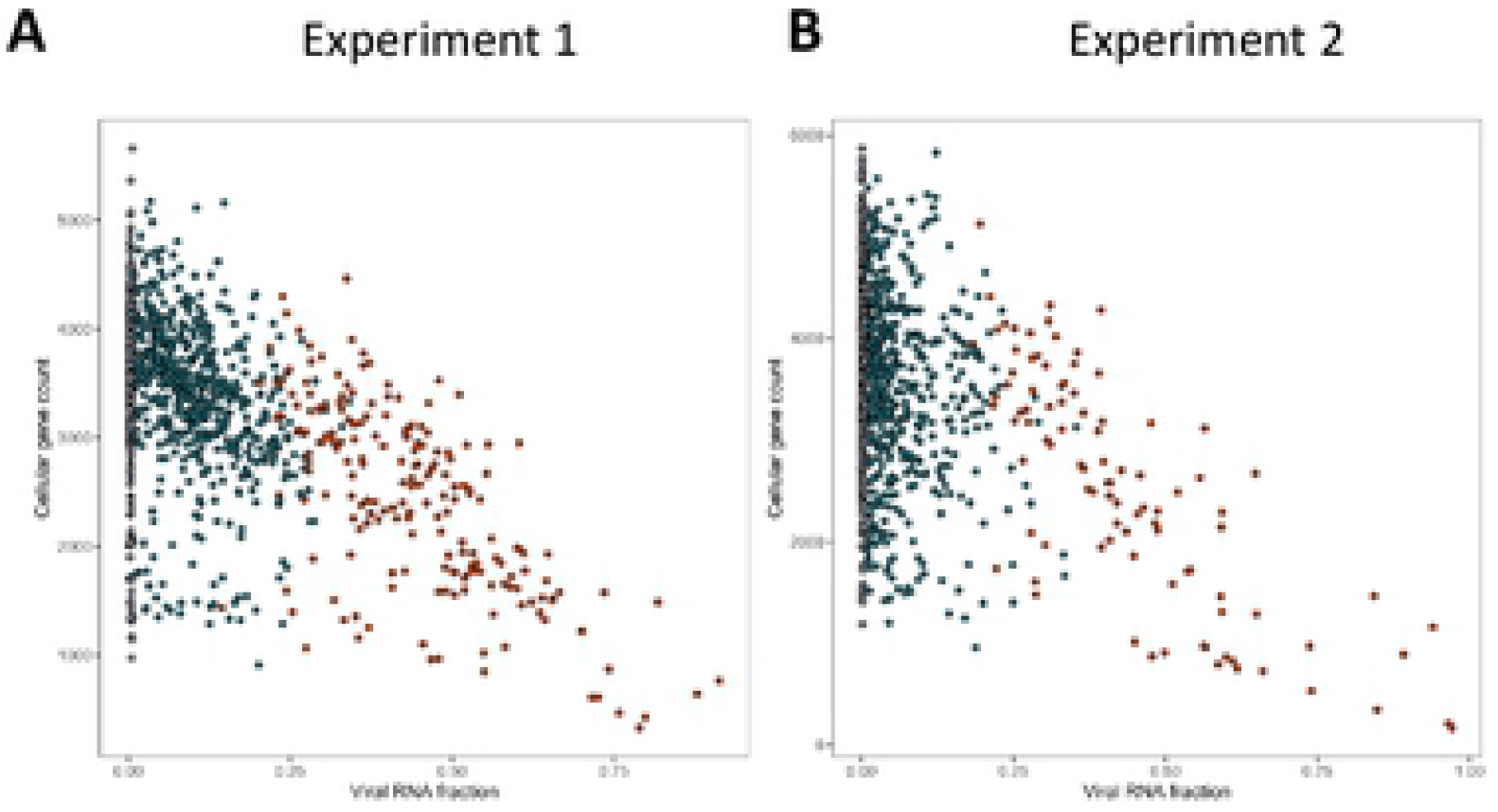
Correlation of cellular and viral genes expression measured by single-cell RNA sequencing. Cells from two independent TNBC were either mock infected or infected with VV (MOI = 5). Six hours later the cells were subjected to the 10X Genomics single-cell protocol, followed by sequencing. Dots represent cells positioned according to the percentage of viral gene expression (x-axis) and the number of cellular genes expressed (y-axis). A: experiment 1; **B**: experiment 2.

### Differential expression analysis using naïve, bystander and infected cells

For the 2 experiments, a standard statistical analysis using Seurat v3 was performed using cells with a percentage of mitochondrial genes below 25%. On the UMAP plots produced, the cells segregated in three (experiment 1, Fig. 5A) and four (experiment 2, Fig. 6A) clusters. These clusters contained both control cells and cells exposed to the virus (Fig. 5B and 6B). Three distinct cellular populations were distinguished among the different clusters: naïve cells, defined as cells not exposed to VV; bystander cells, defined as cells exposed to the virus but expressing less than 0.01 % of early viral genes; infected cells, defined as expressing more than 0.01 % of early viral genes. Naïve, bystander and infected cells were localized onto the UMAP plot (Fig 5C and 6C). The distribution indicated that the clusters showing the higher proportion of bystander cells in the two experiments are the “COL1A2” clusters. The relative proportion of the three subpopulations (naïve, bystander, infected) in all the clusters are presented in Table S1. Assuming that a higher proportion of bystander cell within a cluster is associated with an increased refractoriness to the virus, we looked upstream regulators associated with the two “COL1A2” clusters using Ingenuity Pathway Analysis™ (IPA) analysis. The transcriptomic signatures of the cells show, for the two clusters, a pattern highly consistent with “TGF-β” as a major upstream regulator (Figure S2).

**Figure 5:**
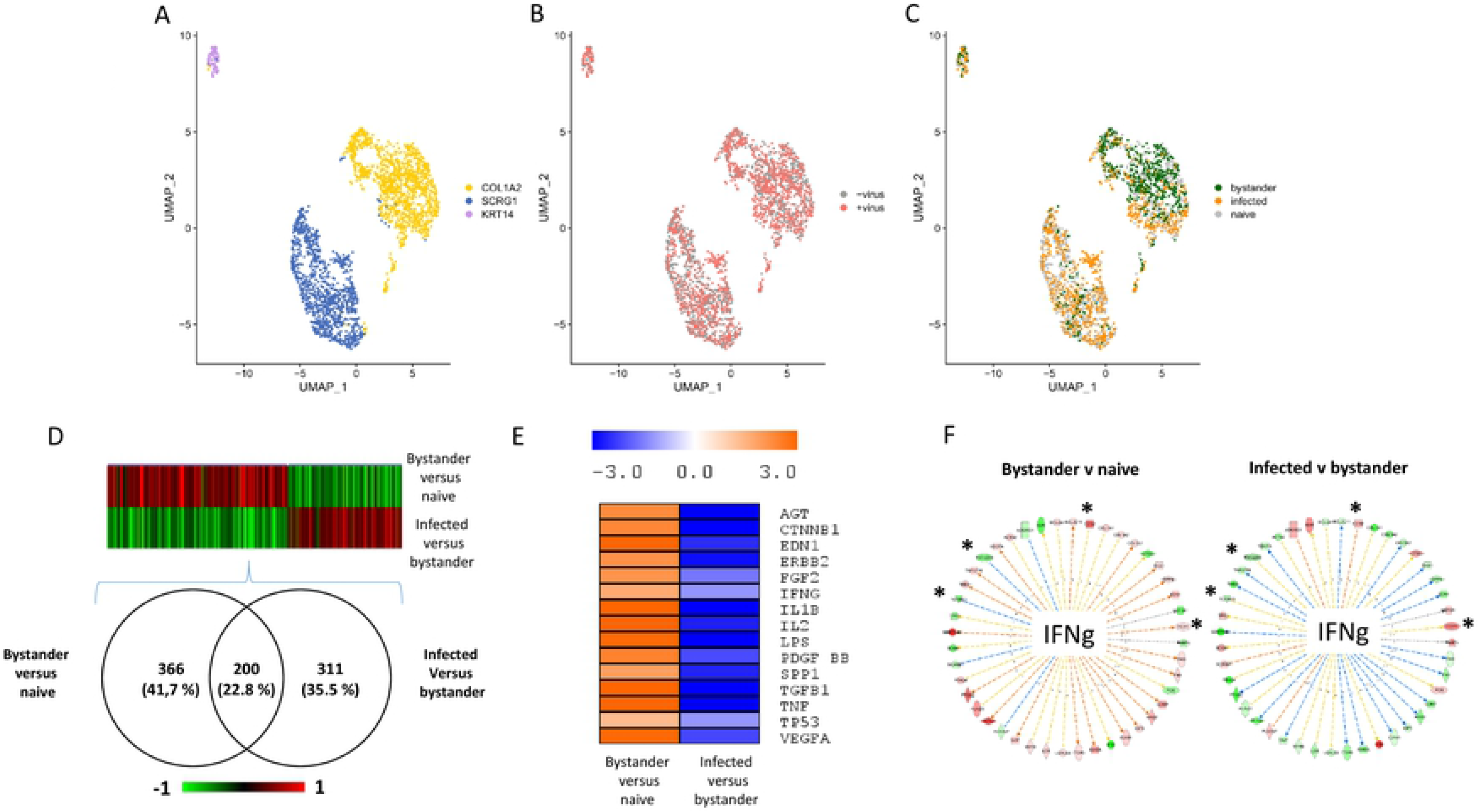
Single-cell transcriptomic analysis of TNBC infected with VV (experiment 1). Cells from TNBC were either mock infected or infected with VV (MOI = 5). Six hours later the cells were subjected to the 10X Genomics single-cell protocol, followed by sequencing. **A**: UMAP representing the three clusters annotated “COL1A2”, “SCRG1” and “KRT14” as these genes are top discriminating of the three clusters. **B**: Repartition of the cells incubated (red) or not (grey) with the virus. **C**: Repartition of naïve, bystander and infected cells in the three clusters. **D**: Venn diagram representing genes that are modulated in bystander versus naïve cells and infected versus bystander cells. The pattern of expression of the genes commonly regulated in the two differential analysis is presented as a heat-map. Red: gene upregulated, green gene down regulated. **E**: Ingenuity Pathways Analysis showing the upstream regulators describing differentially expressed genes in bystander versus naïve cells and infected versus bystander cells. Orange: pathway activated; blue: pathway inhibited. **F**: Example of genes part of the IFNγ pathway inversely regulated in bystander versus naïve cells and infected versus bystander cells. Only four out of 44 genes (9.1 %) are regulated in the same direction in the two conditions. These genes are noted: *.

**Figure 6:**
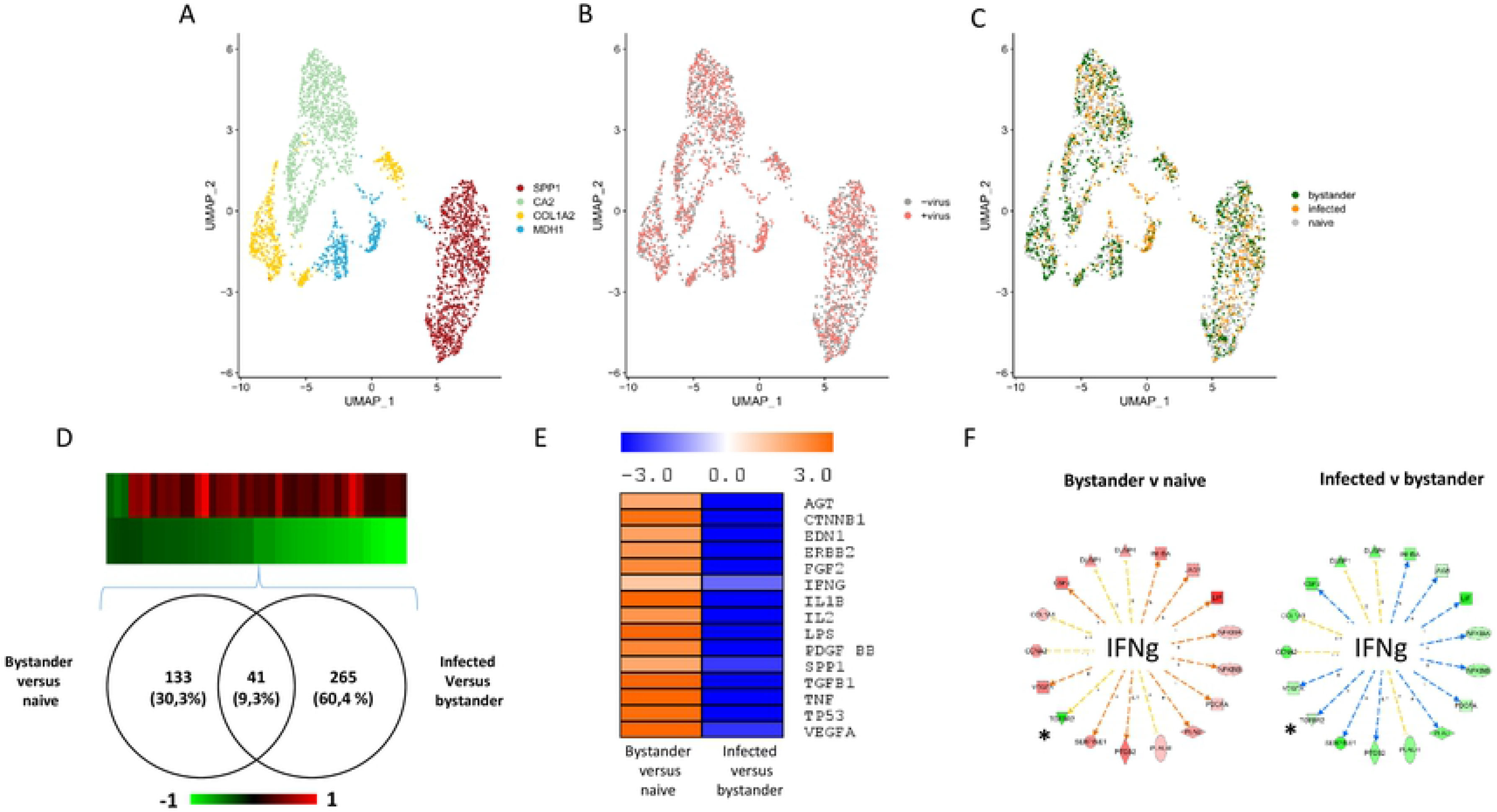
Single-cell transcriptomic analysis of TNBC infected with VV (experiment 2). Cells from TNBC were either mock infected or infected with VV (MOI = 5). Six hours later the cells were subjected to the 10X Genomics single-cell protocol, followed by sequencing. **A**: UMAP plot representing the four clusters annotated “SPP1”, “CA2”, “COL1A2” and “MDH1” as these genes are top discriminating of the four clusters. **B**: Repartition of the cells incubated (red) or not (grey) with the virus. **C**: Repartition of naïve, bystander and infected cells in the four clusters. **D**: Venn diagram representing genes that are modulated in bystander versus naïve cells and infected versus bystander cells. The pattern of expression of the genes commonly regulated in the two differential analysis is presented as a heat-map. Red: gene upregulated, green gene down regulated. **E**: Ingenuity Pathways Analysis showing the upstream regulators describing differentially expressed genes in bystander versus naïve cells and infected versus bystander cells. Orange: pathway activated; blue: pathway inhibited. **F**: Example of genes part of the IFNγ pathway inversely regulated in bystander versus naïve cells and infected versus bystander cells. Only one out of 17 genes (5.9 %) are regulated in the same direction in the two conditions. This genes is noted: *.

To describe the molecular events associated with viral infection, a differential expression analysis was performed between bystander and naïve cells and between infected and bystander cells. The whole dataset is presented in Supplemental Table 2A and B (experiment 1) and Table 3A and B (experiment 2). Figure 5D shows that 200 genes were commonly-regulated in bystander versus naïve and infected versus bystander and an inverse regulation of these commonly-regulated genes was observed (Fig. 5D). IPA analysis of the differentially expressed genes provided information on the upstream regulators describing the differences between bystander and naïve cells and between infected and bystander cells. Figure 5E shows that activation of the pathways regulated by, for example, TGF-β1, TNF, IL1β or IFN-γ can be observed when bystander cells are compared to naive cells. This pattern is likely to reflect the reaction of bystander cells to the presence of the virus in the culture medium and to the secretion of various cytokines by cells infected with VV. By contrast, these pathways were inhibited when the IPA analysis was performed on the differentially expressed genes between infected cells and bystander cells (Figure 5E). A similar phenomenon was observed in the second experiment (Figure 6D and 6E). However, in this second experiment, the number of genes modulated in bystander minus naïve cells was lower than that observed in experiment 1 (41 genes, see Fig. 6D). As the TNBC cells used in experiment 2 are 10 times less sensitive to the virus than those used in experiment 1, these differences may be attributed to a blunted ability of cells more resistant to the virus to respond to the presence of the virus in the culture medium and to stimuli secreted by infected cells. Finally, the striking contrasts between bystander minus naïve cells and infected versus bystander cells were also observed at the level of individual pathways. For example, more than 90% of genes of the IFNγ pathway that were regulated in a particular direction in bystander versus naïve cells were regulated in the opposite direction in the infected versus bystander cells (Fig. 5F and 6F).

### Identification of genes overrepresented in bystander versus infected cells

We hypothesized that genes with “antiviral” activities were overrepresented in the bystander compared to the infected population of cells. Fig. 7A shows the Venn diagram of the genes differentially expressed in bystander cells in experiments 1 and 2. Under our hypothesis, the 130 genes commonly regulated are candidate genes with antiviral activities (complete list in Table S4). IPA analysis showed that these genes were consistent with an activation of the pathways regulated by TGFβ1, LPS, TNF, CTNNB1 and IL1β (Fig.7A), suggesting that activation of these pathways are associated with an antiviral action. The comparison of the 130 candidates with genes identified as potential anti-viral genes in high throughput RNAi screens showed that only one gene, SERPINE1 was found in common with the studies of Beard et al. [12] and none with the study of Sivan et al. [11] (Fig.7B).

**Figure 7:**
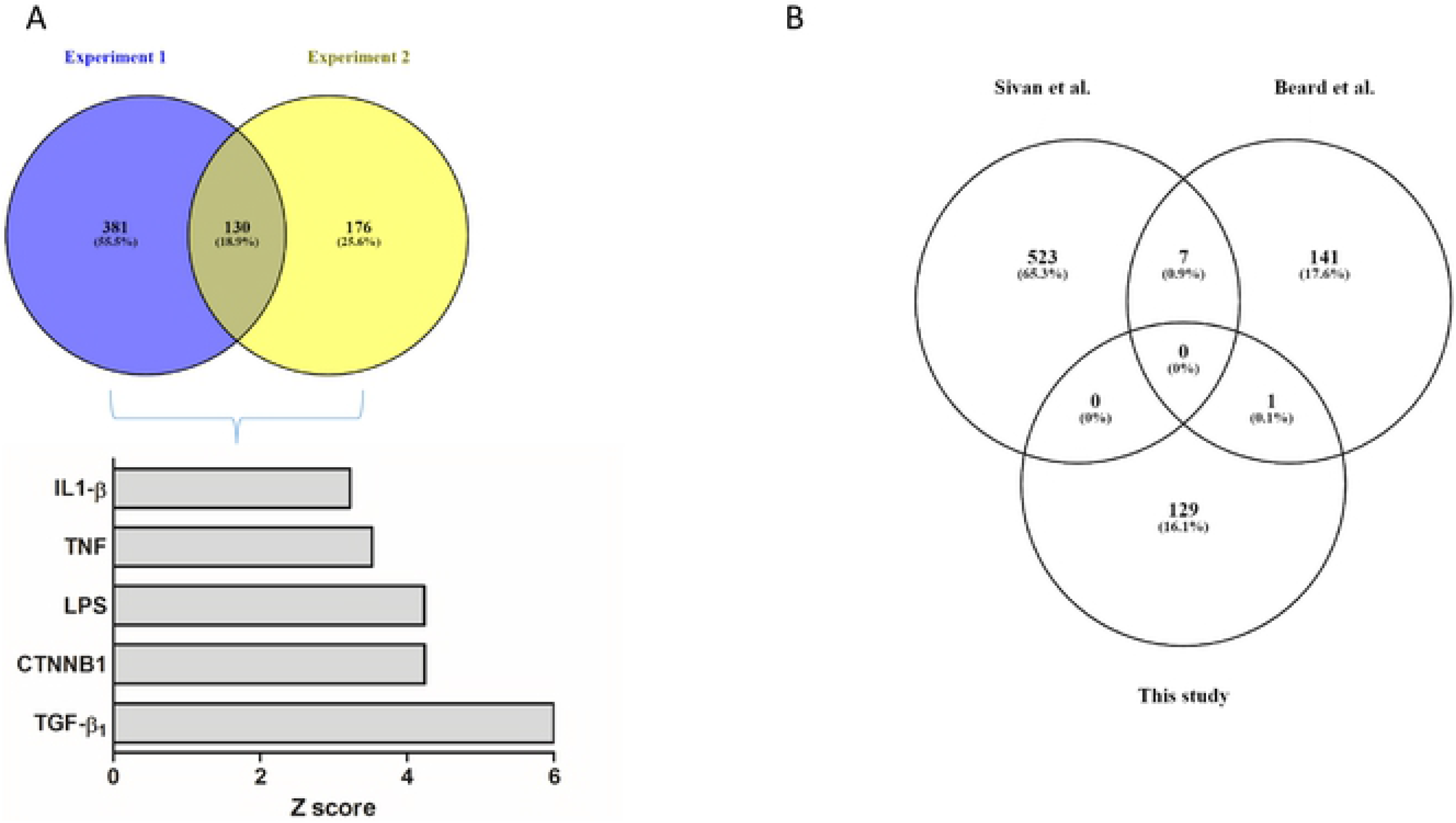
Analysis of the differentially-expressed genes in bystander versus infected cells. **A**: Venn diagram of the differentially-expressed genes in bystander versus infected cells in experiment 1 and 2. A total of 125 genes are commonly regulated. Ingenuity Pathways Analysis showed that these genes were consistent with an activation of the pathways regulated by TGFβ1, CTNNB1, LPS, TNF and IL1β. The z-score for each pathway is presented. **B**: Comparison of the potentially “antiviral genes” in the present study and in the studies of Sivan et al. [11] and Beard et al. [12].

### DDIT4 exerts an antiviral activity

An alternative way to analyze the dataset is to consider each individual clusters in each experiment. This analysis grants less weight to clusters with high number of cells. We used the FindConservedMarkers command in Seurat v3, to run differential expression tests cluster by cluster in order to identify the conserved markers between bystander and infected cells. We required a gene to have a log_2_ (Fold Change) > 0.25, and a maximum Bonferroni-corrected P value threshold < 0.05 to be considered as a conserved marker. This analysis identifies genes that are differentially regulated between two conditions (i.e. bystander versus infected) across all clusters in one experiment. We identified 19 and 79 conserved genes in experiments 1 and 2, respectively. Interestingly, only 7 genes were conserved between the two experiments (Fig.8A). Two of these genes were canine genes for which human homologs have not been identified (ENSCAFG00000032813 and ENSCAF00000031808). The five remaining genes are APEX1, DDIT4, DUSP6, TBCB and DUSP1. The latter has already been shown to be detrimental to vaccinia virus [8].

We focused on one particular gene (DDIT4) in the list presented in Fig. 8A, as high DDIT4 expression has been associated with a poor prognosis in various malignancies that include breast cancer [30] and in TNBC [31]. We therefore examined the effect of DDIT4 on VV replication. Infection of HeLa cells overexpressing DDIT4 resulted in a 60% reduction in the production of infectious VV particles compared to control HeLa cells expressing GFP (Fig. 8B). Inversely, infection of mouse embryonic fibroblast (MEF) from DDIT4 knock out mice resulted in a six-fold increase in the production of infectious VV particles compared to MEF isolated from wild-type mice (Fig. 8C).

**Figure 8:**
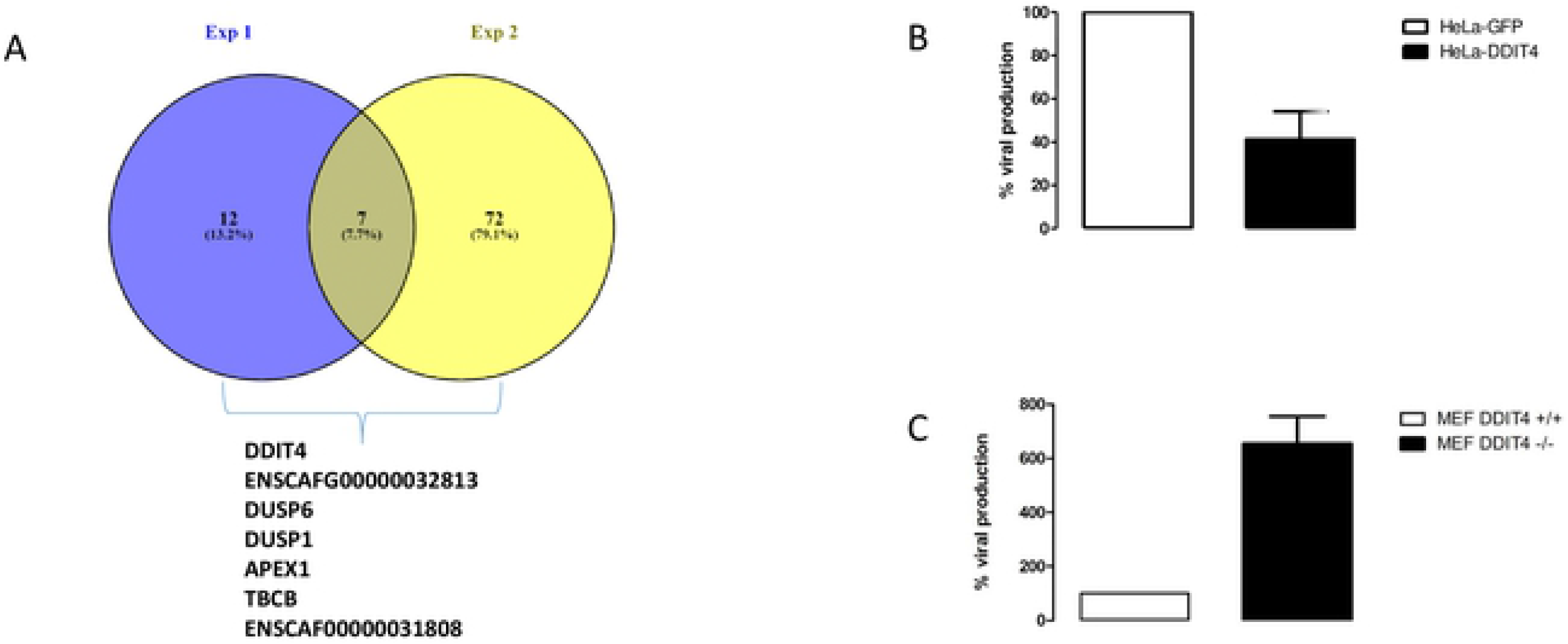
Analysis of the differentially-expressed genes in bystander versus infected cells using the “conserved markers” strategy. **A**: Venn diagram of the differentially-expressed genes in bystander versus infected cells in experiment 1 and 2, using a conserved marker analysis. The 7 commonly regulated genes in the two experiments are indicated. ENSCAFG00000032813 and ENSCAF00000031808 are canine genes without human homologs. **B** Three days after VV infection, HeLa-GFP (white bars) or HeLa-DDIT4 (black bars) cells were collected, homogenized and the number of infectious particles generated in 30 µL of homogenates were titrated. The results are expressed as percentage of infectious particles obtained from HeLa-GFP cells **C**: Three days after VV infection, wilt-type MEF (white bars) or MEF DDIT4^−/−^ (black bars) were collected, homogenized and the number of infectious particles generated in 30 µL of homogenates were titrated. The results are expressed as percentage of infectious particles obtained from wild-type MEFs. For B and C, the results are mean +/− SD of three independent determinations.

## Discussion

We demonstrate that oncolysis induced by VV in primary, high-grade canine mammary carcinoma is significantly less efficient than in equivalent cells obtained from lower grade tumors. This observation is in sharp contrast with the fact the same virus is equally efficient in established cell lines from differentiated/low-grade and in high-grade, human TNBC. Considering the close relationship between the human and canine pathologies [32], it is tempting to attribute this difference in effectiveness of VV to the primary/low passage versus established cell lines status of the experimental models. The relevance of established cell lines as experimental systems to develop new cancer therapeutic agents has largely been questioned in the past and the need for new preclinical models has been highlighted [33]. Patient-derived xenografts have been proposed and are viewed as one of the most relevant model systems in oncology [34]. For a selected number of types of tumors that include breast cancer, canine tumors recapitulate the features of human ones and resources from relevant canine tumors have been proposed as tools for the preclinical development of new cancer therapeutics [35]. One of these resources is very low passage primary cells grown in serum-free medium. Working with primary, low-passage cells has been to date hampered by the small number of cells available from biopsies. It is rare to collect more than 2-3 million carcinoma cells from one biopsy, and without amplification, this low number of cells restricts considerably the information that can be gathered experimentally. However, with the advent of single-cell transcriptomics, descriptive studies demonstrating whether a therapeutic agent is effective or not can be complemented with high-resolution molecular data.

Single cell transcriptomic has been used previously in the field of infectious diseases. For example, the extreme heterogeneity of influenza virus infection [36] and influenza infection of mouse lungs in vivo [37] have been examined with this tool. But, to our knowledge, it has never been applied to the study of poxvirus infection. First, our study confirms the well-documented transcriptional shut-down of cellular genes. In our dataset, this shut-down is correlated to the extent of viral gene expression (Figure 4). A unique feature of single-cell transcritptomic analysis is the possibility of dissecting different populations of cells that have been in contact with the virus. Cells expressing intermediate and late viral genes express a low number of cellular genes. Their inclusion in the analysis did not provide any particular information for identification of antiviral genes. By contrast, bystander cells (cells exposed to the virus and expressing less than 0.01 % of viral genes) provide a unique source of information. Comparison of bystander and naïve cells showed, an activation of the pathways regulated by cytokines and growth factors in bystander cells. These activations are likely to be the results of the combined action of the pathogen-associated molecular pattern of the virus as well as autocrine factors secreted by infected and dying cells. The comparison of the activations observed in the two single-cell transcriptomic experiments shows that in the second experiment that involves a culture of TNBC cells more refractory to the virus, a blunted response is observed, that involves the number of genes and the degree of modulation of the regulated genes. Inversely, an inhibition of cytokine and growth factor pathways can be observed when infected versus bystander cells are compared. These inhibitions are likely to result from the expression of viral genes that counteract the cellular responses.

Our study demonstrates that the information gathered through single-cell RNA-seq can lead to the identification of pathways with potential antiviral properties. We hypothesized that genes with “antiviral” activities were overrepresented in the bystander compared to the infected population of cells. Differential expression analysis on the bulk set of data identified 130 genes commonly overrepresented in the bystander populations in the two experiments performed. IPA analysis identified upstream regulators likely to produce these expression profiles (Fig 7A). The constitutive activation of the interferon pathway has been already documented as an “anti-oncolytic” mechanism for different viruses [38, 39], and this pathway, as well as general pathways associated with inflammatory responses (IL1β, lipopolysaccharides (LPS) TNF) are also identified in our dataset with poxvirus infection (Fig7A). Additional pathways appear to characterize cells from the bystander population: CTNNB1 (β-catenin) and TGF-β (Fig.7A), the latter having been already identified as a characteristic for the COL1A2 cluster showing increased “resistance” to the virus (Fig.5A and 5C). These two pathways, associated with the “inflammatory” pathways have been largely implicated in the epithelial to mesenchymal transition (EMT) in general and in EMT in breast cancer in particular [40–42]. It is therefore tempting to postulate that cells with a higher “EMT index” would be less sensitive to vaccinia virus. This hypothesis will have to be thoroughly tested in future studies.

A list of 130 candidates is presented in Supplemental Table 4. The comparison with genes identified as potential anti-viral genes in high throughput RNAi screens showed that only one gene (SERPINE1) was found in common with these studies (Fig. 7B). This low overlap is hardly surprising considering that both virus and cells are different in these screens. Nevertheless, it highlights the complexity of the interactions of vaccinia virus with host cells.

An analysis taking into account the individual clusters provided a more stringent test, with only seven genes emerging as commonly overrepresented in bystander cells, in the two experiments (Fig.8A). Two of those are canine genes with no known human homolog. The five remaining genes are APEX1, DDIT4, DUSP1, DUSP6 and TBCB. The fact that DUSP1 expression has already been demonstrated to be detrimental to the virus [8] provides a reinsurance on the validity of the hypothesis whereby “antiviral” genes were overexpressed in the bystander populations and highlights the relevance of the methodology we used. The proteins encoded by DUSP1 and DUSP6 are phosphatases with dual specificity for tyrosine and serine. They can dephosphorylate MAPK1/ERK2. DUSP1 has been shown to be involved in the replication and host range of vaccinia virus and in the regulation of host immune responses through the modulation of MAPKs [8]. Although DUSP1 and DUSP6 have been attributed different roles, in immune regulation and development, respectively [43], they may exert antiviral activities through similar mechanisms. Tubulin folding cofactor B (TBCB) has been shown to be required for microtubule network formation [44] but, to our knowledge, an involvement of TBCB in any viral infection has never been shown. However, considering the major cytoplasmic reshuffle observed in vaccinia-virus-infected cells, a negative role of TBCB would not be surprising. APEX1 is a major apurinic/apyrimidic (AP) endonuclease in human cells. AP sites are pre-mutagenic lesions and this enzyme is therefore part of the DNA repair machinery [45]. Relationships between APEX1 and viral infection have been documented: inhibition of APEX1 redox activity affects Kaposi’s sarcoma-associated herpes virus [46] and its knock-down inhibits HIV1 and HIV2/SIV infection [47]. However, a role of APEX1 in vaccinia virus lytic cycle has never been reported.

We decided to investigate whether DNA damage inducible transcript 4 (DDIT4) affects VV replication and we show, using gain- and loss-of function that this is the case (Fig.8B and 8C). DDIT4 is expressed in breast cancer and is even associated with a poor prognosis in various cancers that include breast cancers [30]. In high-grade, triple-negative breast cancers, DDIT4 is also associated with a poor prognosis in human patients [31]. This observation positions DDIT4 as a potential marker that may also be associated with a lower response to oncolytic vaccinia virus. Considering the association of DDIT4 with a worse prognostic in human patients with acute myeloid leukemia, glioblastoma multiform, colon, skin and lung cancers in addition to breast cancer[30], future preclinical and clinical studies will determine the real importance of this gene in the response of various cancer types to oncolytic VV. DDIT4 is an interferon-stimulated gene with anti-retrovial activity [48]. Biochemically, DDIT4 has largely been described as a negative regulator of the mTOR signaling pathway [49–51]. Rapamycin, a pharmacological inhibitor of the mTOR signaling pathway, has also been described to reduce the virus yield upon VV infection [52]. A possible mechanism may be that mTOR activation results in the phosphorylation of 4E-BP, which in turn releases the translation factor elF4E, the component of el4F that binds to the 5’-cap structure of mRNA and promotes translation [52, 53]. Upon VV infection, the factor elF4E has been reported to be redistributed in cavities present within viral factories [27, 54] where viral translation can proceed. It is therefore tempting to hypothesize that DDIT4, by inhibiting the mTOR signaling pathway, reduces the amount of elF4E available for viral translation. However, considering the complexity of mTOR effects on VV infection [55], the exact nature of the molecular events associated with the inhibitory effect of DDIT4 remains to be elucidated. Finally, the identification of cellular genes promoting or restricting vaccinia virus infectivity/replication has been studied using hypothesis-driven approaches [8, 9, 56] or high-throughput RNA interference screens [10–13] and in this context single-cell transcriptomics increase the arsenal of experimental tools available. Considering the large transgene capacity of poxviruses, this information could be exploited to generate new generations of oncolytic poxviruses with more efficient direct oncolytic properties.

## Materials and Methods

### Cells

Very low passage canine primary cell cultures were provided by Lucioles Consulting. They were derived from a panel of canine primary tissues including normal mammary tissues, hyperplastic lesions, benign tumors, carcinomas in situ and all grades of carcinomas. A list of the different biopsies is provided in Table S5. The tissues were phenotyped using standard histopathology and immunohistochemistry techniques. Cell survival assays were performed as previously described [57]. BHK21, MCF7, MDA-MB231, HeLa and DDIT4 +/+ and −/− MEF cells were obtained and cultured as previously described [50, 58–62]. Fluorescence imaging and Western blots were performed as previously described [63, 64], respectively. Quantification of fluorescent cells were performed using the CellQuant program (available at: http://biophytiro.unice.fr/cellQuant/index_html).

### Viruses

A VACV-Lister strain deleted in the thymidine kinase gene (referred to as VV) and VACV-Copenhagen recombinants encoding GFP downstream of a synthetic early promoter (VACV-Cop21 and VACV-Cop 32) were described previously [65, 66]. Vaccinia virus titration was performed on BHK21 cell monolayers infected for two days and stained with neutral red. Lentiviruses (encoding either GFP or DDIT4) were purchased from Sigma-Aldrich and are part of the MISSION TRC3 LentiORF collection.

### qPCR assay for generic detection of Orthopoxvirus

DNA was extracted using the QIAamp DNA Mini Blood kit (QIAgen). The qPCR assay used for the detection of orthopoxviruses was a modification of the assay described by Scaramozzino et al [67]. Probes were designed to amplify a 157 bp fragment of orthopoxvirus A27L gene. Each qPCR assay was carried out in 20µl of a reaction mixture containing 5µl of extracted DNA as template, 400 nM of each primer and 250 nM of probe and 10 µl of IQ Supermixe for QPCR (Biorad). The reaction was performed as follows: 1 cycle at 95°C for 3 min, followed by 45 cycles each at 95°C for 15s, followed by 62°C for 60 s. A fluorescence reading was taken at the end of each 62°C step. Data acquisition and analysis were carried out with the Bio-Rad CFX Manager software 3.1. Sample curves were analyzed by using the second derivative. Each DNA solution was assayed in duplicate per qPCR assay. Standard curves were generated from serial dilution of a solution of pVACV_Lis-A27L enabling absolute quantification.

### qPCR assay for generic detection of early and late genes, E9L and A27L respectively, and a housekeeping gene

Total RNA was extracted using the RNAeasy kit (Qiagen). Reverse-transcription was performed using the PrimeScript™ RT Reagent Kit (Perfect Real Time, TAKARA). The A27L (late gene) qPCR assay was performed as described above. The E9L (early gene) and beta-actin qPCR were performed as described by Kulesh et al.[68] and Piorkowski et al. [69], respectively, with slight modifications. Each qPCR assay was carried out in 20µl of a reaction mixture containing 5µl of cDNA as template, 300 nM of each primer and 100 nM of probe and 10 µl of IQ Supermixe for qPCR (Biorad). Each DNA solution was assayed in duplicate per qPCR assay. Standard curves were generated from serial dilution of a CPXV supernatant. PCR efficiencies of both the targets genes and the reference gene were between 90 and 110% and did not differ by more than 10%. The delta Ct method was then used for relative quantification.

### Bulk RNA-sequencing

Cells were either mock infected or infected at a MOI of 5 with VV. At different time after infection, cells were washed with PBS. Poly(A) RNAs were purified using a Dynabeads mRNA purification kit (Invitrogen) and fragmented for 7 min at 95 °C. Libraries were then generated with the Ion Total RNA seq kit V2 (Life technologies) and sequenced on the Ion Proton system with P1 chip V3 following the manufacturer’s instructions. Reads were aligned to the dog genome release canFam3 and the Vaccinia virus genome release NC006998 withbowtie v2-2.2.4. Quantification of genes was then performed using HTSeq-count release HTSeq-0.6.1 with “--minaqual=0--mod=intersection-nonempty” options. To assess the response differences to the viral infection between Non TNBC and TNBC samples, we used the classification of early, intermediate and late poxvirus genes described previously reported [70]. P-values on boxplots were calculated by the Wilcoxon rank sum test.

### Single Cell RNA-sequencing

Single-cell suspensions were converted to barcoded scRNA-seq libraries by using the Chromium Single Cell 3’ Library, Gel Bead & Multiplex Kit and Chip Kit (10x Genomics), aiming for an estimated 2,000 cells per library and following the manufacturer’s instructions. Samples were processed using kits pertaining to V2 barcoding chemistry of 10x Genomics. Libraries were sequenced on an Illumina NextSeq500 with a High Output v2 kit (150 cycles): the forward read had a length of 26 bases that included the cell barcode and the UMI; the reverse read had a length of 98 bases that contained the cDNA insert. Raw sequencing FASTQ files were analyzed within 10x Genomics CellRanger suite (v1.3.0) with a transcriptome reference composed of canFam3 Canis familiaris genome build and the Vaccinia virus complete genome (NCBI reference sequence NC_006998).

### Single-cell gene expression quantification and determination of the major cell types

Raw gene expression matrices generated with 10xGenomics CellRanger suite (v1.3.0) were loaded and processed into R (version 3.5.2). Both experiments were analyzed independently using the Seurat R package (version 3.0.0). First, all cells that had over 25% of mitochondrial RNAs were removed. From the remaining cells, gene expression matrices were normalized using SCTransform method. To reduce dimensionality, variably expressed genes were summarized by principle component analysis, and the 30 PCs were further summarized using UMAP dimensionality reduction. Both samples (i.e. infected and not infected) from the two experiments (i.e. 1 and 2) were then aggregated using FindIntegrationAnchors and IntegrateData functions preceded by the PrepSCTIntegration function described in Seurat development version. Clusters were called using a low resolution of 0.1, and gene markers were assessed using FindAllMarkers function with standard parameters.

## Data and code availability

Transcriptomics single cell and bulk data can be accessed on NCBI GEO accession number GSE142185. R Code and preprocessed objects required to reproduce analysis can be accessed on github (https://github.com/ucagenomix/sc.cambien.2020).

## Acknowledgements

This work was performed with the financial support from ITMO Cancer AVIESAN (Alliance Nationale pour les Sciences de la Vie et de la Santé, National Alliance for Life Sciences & Health) within the framework of the Cancer Plan (“Plan Cancer 2009-2013”, “Plan Cancer 2014-2019”) and with support from the Canceropole PACA. Special thanks are due to Dr Robert Drillien (Strasbourg, France) for providing viruses, help and protocols in vaccinia handling and careful reading of the manuscript.

